# Deep learning models for unbiased sequence-based PPI prediction plateau at an accuracy of 0.65

**DOI:** 10.1101/2025.01.23.634454

**Authors:** Timo Reim, Anne Hartebrodt, David B. Blumenthal, Judith Bernett, Markus List

**Author notes:** Corresponding and joint last authors.

## Abstract

As most proteins interact with other proteins to perform their respective functions, methods to computationally predict these interactions have been developed. However, flawed evaluation schemes and data leakage in test sets have obscured the fact that sequence-based protein-protein interaction (PPI) prediction is still an open problem. Recently, methods achieving better-than-random performance on leakage-free PPI data have been proposed. Here, we show that the use of ESM-2 protein embeddings explains this performance gain irrespective of model architecture. We compared the performance of models with varying complexity, per-protein, and per-token embeddings, as well as the influence of self- or cross-attention, where all models plateaued at an accuracy of 0.65. Moreover, we show that the tested sequence-based models cannot implicitly learn a contact map as an intermediate layer. These results imply that other input types, such as structure, might be necessary for producing reliable PPI predictions.

## 1. Introduction

Proteins perform a wide array of biological functions, but more than 80% depend on protein-protein interactions (PPIs) [Zhou et al., 2016]. A comprehensive understanding of PPIs could significantly improve our knowledge of biological mechanisms, facilitate the identification of therapeutic targets, and aid in designing drugs for modulating specific protein interactions [Rao et al., 2014]. Numerous experimental techniques, such as yeast two-hybrid, tandem affinity purification-mass spectrometry, X-ray crystallography, and NMR spectroscopy, have been developed to identify PPIs [Howell et al., 2006, Rao et al., 2014]. However, they are cost-intensive and low-throughput compared to computational approaches [Rao et al., 2014]. Since PPIs inherently occur in a three-dimensional context, including structural data would be ideal for this task. Unfortunately, obtaining 3D structures remains challenging despite the availability of in silico prediction tools like AlphaFold [Jumper et al., 2021]. Hence, predicting PPIs based solely on amino acid sequences, which are more accessible, could provide an efficient alternative.

In a recent study, we highlighted issues in sequence-based PPI prediction methods, where improper data splitting introduced data leakage, inflating reported accuracies to levels exceeding 90% [Bernett et al., 2024]. The leakage resulted from sequence similarity and protein node degree shortcuts between the training and testing sets. Re-evaluating these models on a leakage-free gold-standard dataset [Bernett, 2022] revealed significantly lower and often close to random performances, emphasizing the need for careful dataset preparation for unbiased method evaluation.

Since the identification of these issues, two notable models have been developed using our gold-standard dataset. Sledzieski et al. [2024] employed parameter-efficient fine-tuning (PEFT) of Evolutionary Scale Modeling-2 (ESM-2) embeddings. They achieved the best results with a baseline fully-connected neural network (FCNN) (accuracy around 0.63) from which they concluded that the lack of large datasets is currently the bottleneck in PPI prediction. Further, they found that training the FCNN on the smallest version of the ESM-2 embeddings (t33: 650 million parameters) yielded better results than the larger embeddings. Another model, TUnA, introduced by Ko et al. [2024a], applies transformer encoders followed by spectral normalized neural Gaussian processes (SNGPs) for uncertainty-aware predictions. This architecture integrates spectral normalization with a Gaussian process to improve uncertainty estimation. They reached an accuracy of approximately 0.65. Additionally, three studies were published, enhancing the ESM-2 protein embeddings and evaluating their performance on our gold-standard dataset using only simple ML baselines. ProteinCLIP [Wu et al., 2024] combines ESM-2 and ProtT5 per-protein embeddings with natural language embeddings of protein function by OpenAI’s text-embedding-3-large model. They achieve 0.61 - 0.65 accuracy. NaderiAlizadeh and Singh [2024] argue that dimension-wise average pooling to go from a per-residue to a per-protein representation is suboptimal as positions should not be weighted equally. They introduce a new pooling approach based on optimal transport. With both the ESM-2 average pooling approach and their new approach, they achieve about 0.65 accuracy. PoolPaRTI [Tartici et al., 2024] computes per-protein embeddings by combining the internal attention matrices of the ESM-2 model using PageRank and also reach about 0.65 accuracy.

The fact that all of these approaches reach similar accuracies of around 0.65 with very different architectures and complexity levels raises a simple question: Are models for unbiased sequence-based PPI prediction attainable that yield accuracies that are substantially better than 0.65? Or is it likely that the field will plateau at this performance level, which would imply that focusing on models relying on other input types (e.g., structures) is necessary for reliably predicting PPIs?

In this work, we sought to answer this question by systematically assessing the effect of various techniques used in recently proposed PPI prediction models. Specifically, the state-of-the-art protein language model (PLM) embeddings ESM-2 [Lin et al., 2022], as well as self and cross-attention modules, were assessed to gauge their individual and combined contribution to prediction performance. Previously published models such as Richoux [Richoux et al., 2019] and D-SCRIPT [Sledzieski et al., 2021] were modified using ESM-2 embeddings which led to notable improvements. The impact of model architecture and complexity was analyzed, comparing FCNNs, convolutional neural networks (CNNs), and transformer encoder-based models. Results indicate that embedding quality plays a more critical role in performance than model architecture, as no model achieved an accuracy higher than 0.65. Lastly, the possibility of implicitly predicting contact maps of interacting proteins as a penultimate layer was also investigated to assess its validity, as this approach has been previously suggested [Sledzieski et al., 2021].

## 2. Methods

### 2.1. Data

Our previously introduced gold-standard dataset [Bernett, 2022] was used for training, validation, and testing. This dataset is divided into three groups, Intra0, Intra1, and Intra2, with 59 260, 163 192, and 52 048 entries, respectively, each with an equal number of positive and negative interactions. The positive interactions stem from the HIPPIE v2.3 database [Alanis-Lobato et al., 2017]. Intra1 is used for training, Intra0 for validation, and Intra2 for testing. This dataset is leakage-free since no protein is present in more than one split and sequence similarity between the splits is minimized by construction (we used KaHIP [Sanders and Schulz, 2013] to partition proteins based on sequence similarity and further reduced redundancy in with CD-HIT at 40% pairwise sequence similarity). With proteins having approximately the same node degree in the positive as in the negative dataset, node degree bias is also strongly reduced.

### 2.2. Protein embeddings

We generated protein embeddings from the amino acid sequences using ESM-2 [Lin et al., 2022], which then served as the input for the models. Embeddings from the three largest ESM-2 models, with dimensions *d*_*emb*_ of 5120, 2560, and 1280 were employed (referred to as t48, t36, and t33 with 15 billion, 3 billion, and 650 million parameters, each). The per-protein embeddings were obtained by dimension-wise average pooling of the per-token embeddings, resulting in a single vector of size *d*_*emb*_ for each protein. For models requiring per-token embeddings, proteins longer than 1000 amino acids were excluded due to computational limitations, leaving 93 719, 46 421, and 41 100 samples in the training, validation, and test sets, respectively.

### 2.3. Models

If not specified otherwise, the training set is always used for training, the validation set for hyperparameter tuning, and the test set for the final evaluation of the models. To counteract overfitting, a random subset containing 50% of the samples of the original training set is used to train the models in each epoch. All deep learning models use binary cross entropy (BCE) as the loss function and Adam for optimization. For models using unpadded per-token embeddings, each sample was processed separately, but backpropagation was performed for the entire batch simultaneously. We reimplemented the D-SCRIPT, Richoux, and TUnA models according to published method descriptions and modifications of the available code that were necessary to adapt the methods for training. Accordingly, even the vanilla reimplementations of D-SCRIPT, Richoux, and TuNA without algorithmic modifications may slightly differ from the model implementations underlying the original publications. Nonetheless, our implementation of Richoux and D-SCRIPT achieved approximately the same results (0.53 vs. 0.52 and 0.5, respectively, Figure S5) as in our previous study when supplied with their original input (i.e., one-hot encoding for Richoux and an embedding from a smaller language model for D-SCRIPT). We only slightly modified TuNA and reached the same accuracy as the original publication (Table 1).

**Table 1.** Comparison of all tested models. Accuracy, precision, recall, F1 score, as well as the binary cross entropy values summarized in the “Loss” column were computed on the test sets. The “Best epoch” column describes the epoch during training with the best accuracy on the validation set, which is 8 epochs before training was terminated due to early stopping. Total training times and mean training times per epoch are reported in seconds.

| Model | Accuracy | Precision | Recall | F1 | Loss | Training time | Best epoch | Training time per epoch |
| --- | --- | --- | --- | --- | --- | --- | --- | --- |
| Baseline models |  |  |  |  |  |  |  |  |
| RFC-40 | 0.577 | 0.629 | 0.374 | 0.496 | – | 474 | – | – |
| 2d-baseline | 0.575 | 0.630 | 0.373 | 0.468 | 0.738 | 30845 | 22 | 1234 |
| Attention models |  |  |  |  |  |  |  |  |
| 2d-Crossattention | 0.641 | 0.660 | 0.589 | 0.623 | 0.643 | 29411 | 6 | 2101 |
| 2d-Selfattention | 0.616 | 0.611 | 0.553 | 0.591 | 0.670 | 41326 | 14 | 1878 |
| Adaptions of published models |  |  |  |  |  |  |  |  |
| Richoux-ESM-2 | 0.633 | 0.627 | **0.654** | **0.640** | 0.653 | 2483 | 8 | **155** |
| D-SCRIPT-ESM-2 | 0.628 | 0.638 | 0.602 | 0.619 | 0.650 | 29849 | 7 | 1990 |
| TUnA | **0.645** | **0.672** | 0.580 | 0.622 | **0.630** | 30013 | 7 | 2001 |

#### 2.3.1 Baseline models

We created two baseline methods for comparison, one each for per-token and per-protein mean embeddings. The baseline models serve as a reference point for assessing the performance of the more complex models. They provide the minimal expectation or basic benchmark that any complex model should surpass to be considered effective. For this reason, hyperparameter optimization was not performed on the baselines.

##### RFC baseline

A random forest classifier (RFC) provides a simple baseline for the one-dimensional per-protein embeddings. The RFC constructs 100 trees, each receiving a random subset of 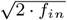 features, with *f*_*in*_ being the total number of input features from one protein. The RFC was trained on the entire concatenated embeddings of both proteins as input. We also tested how much information can be extracted from a dimensionality-reduced ESM-2 embedding. For this, we used the first 200 or 20 components of a principal component analysis (PCA) of the embeddings. The RFC was trained on the concatenated vectors with either 400 or 40 features. The PCA was applied to the entire dataset of mean embeddings. This was done for each of the three embedding sizes. We refer to these models as “RFC-400” and “RFC-40”. We term the unreduced version “RFC-mean”.

##### 2d-baseline

Creating a simple baseline model with the two-dimensional per-token embeddings as input is not trivial, as the variable input size prevents using simple machine learning methods such as RFCs or support vector machines. As a result, the 2d-baseline (Figure S3) is a more complex model but still simpler than the previously published models. It receives the input embeddings of both proteins of dimensions (*len*(*p*_1_), *d*_*emb*_) and (*len*(*p*_2_), *d*_*emb*_) and reduces the embedding dimension with three linear layers with rectified linear units (ReLU) activation functions. The sizes of these layers are *d*_*emb*_*/*2, *d*_*emb*_*/*4 and 64. Next, the outer product along the second dimension of both tensors is computed, resulting in a tensor of shape (*len*(*p*_1_), *len*(*p*_2_), 64). A convolution with one output channel is applied to this, followed by a pooling operation. The maximum value of the resulting tensor is then transformed by the sigmoid function to form the final predicted label.

#### 2.3.2 Attention models

To test the impact of self- or cross-attention, we created modified versions of the 2d-baseline (Figure S4). We inserted an encoder layer between the embedding reduction via linear layers and the outer product of both proteins and applied spectral normalization to all linear layers of the encoder. Both protein embeddings are fed separately through the same encoder. The self-attention model captures within-protein relationships; the cross-attention model focuses on identifying connections between the two proteins. We alse tested a version where the Transformer encoder was applied before the embedding reduction. We call these models “2d-Selfattention”, “2d-Crossattention”, “2d-Selfattention-encoder-pre-reduction”, and “2d-Crossattention-encoder-pre-reduction”. Models using encoder attention without spectral normalization have the suffix “no-spectral”.

#### 2.3.3 Adaptions of published models

For the previously published models, hyperparameter optimization was performed using Weights & Biases (wandb) [Biewald, 2020] with Bayesian optimization of the accuracy on the validation dataset. Optimizations were performed with different parameter combinations for each model, stopping the training process if the validation accuracy did not improve for eight epochs. All per-token models used exclusively the t33 embeddings of size 1280 due to computational limitations. The learning rate and the size of the first linear layer, which reduces the embedding size, were optimized for all models. If present, the dropout rate was also optimized. For models using convolutions and pooling, we optimized the kernel size and pooling type, while the number of heads for multi-head attention and the dimension of the FCNN in the encoder were optimized in the attention models. The exact hyperparameters and their associated range of values can be found in the code.

##### Adaptions of the Richoux model

The model proposed by Richoux et al. [2019] (Figure S1b) was one of the models with a near-random performance on the leakage-free gold-standard dataset introduced in our previous study, making it well-suited to show the impact of the ESM-2 embeddings and other changes. In contrast to the original Richoux model, our adaption receives the per-protein embeddings from each protein as input. The embedding dimensions of each protein are reduced separately by two distinct linear layers. After concatenation, the two resulting vectors are processed through two additional linear layers. The size of the final layer is reduced to 1. From this single value, the predicted label is obtained from a sigmoid activation function. Every layer is followed by ReLU and batch normalization. The model was re-implemented with PyTorch instead of Tensorflow and we refer to it as ‘Richoux-ESM-2”. We also tested a version of the model in which the embeddings are first fed into a Transformer encoder, with and without adding spectral normalization (“Richoux-ESM-2-encoder(-no)-spectral”). Additionally, we tested applying spectral normalization after every linear layer (“Richoux-ESM-2-spectral”).

##### Adaptions of the D-SCRIPT model

The D-SCRIPT model by Sledzieski et al. [2021] was also tested in our previous study and including it here serves a similar comparative purpose as the Richoux-ESM-2 model. We re-implemented D-SCRIPT with PyTorch to ensure consistent testing (“D-SCRIPT-ESM-2”). It takes the variable-size per-token embeddings as input. The model (Figure S1a) reduces the embedding dimension of the per-token embeddings separately for each protein to size *d*, followed by a ReLU function and dropout layer. Next, it computes the absolute difference and element-wise product between each pair of elements in both transformed tensors. These are then concatenated along the last dimension, forming a new tensor of shape (*len*(*p*_1_), *len*(*p*_2_), 2*d*). Using a convolution followed by batch normalization and ReLU, the embedding dimension is reduced to *h*. A matrix of shape (*len*(*p*_1_), *len*(*p*_2_)) is computed with another convolution, batch normalization and ReLU. Sledzieski et al. treat this matrix as an implicitly predicted contact map of the two proteins. Finally, a custom interaction module developed by Sledzieski et al. is applied to generate the predicted label. In short, the interaction module applies a weighting to the matrix, performs max pooling, normalizes the result, and then uses a custom sigmoid function based on non-zero elements in the matrix. We also tested alternative D-SCRIPT adaptions, which included the insertion of a Transformer encoder before or after dimensionality reduction (“D-SCRIPT-ESM-2-encoder-pre-reduction”, “D-SCRIPT-ESM-2-encoder-post-reduction”). We also tested removing the spectral normalization after the encoder (“D-SCRIPT-ESM-2-encoder-pre-reduction-no-spectral”) and applying cross-instead of self-attention in the encoder (“D-SCRIPT-ESM-2-encoder-cross”). The family of D-SCRIPT adaptions is called “D-SCRIPT-like”.

##### Adaptions of the TUnA model

TUnA was developed Ko et al. [2024a] and already uses ESM-2 embeddings as protein representation in the original version. In this model (Figure S2), all input embeddings of proteins with length *<* 1000 are padded to size (1000, *d*_*emb*_). The embedding dimension is first reduced by a linear layer to *d*_*att*_ before the Intra Encoder module is applied. This module utilizes self-attention on the single sequences to recognize important relationships of different positions in the same protein. Though both inputs are processed individually, the weights are shared. The updated tensors *x*1 and *x*2 are concatenated to a combined tensor *x*12, which is then sent to the Inter Encoder module to capture important relationships between the two proteins with self-attention. To ensure permutation invariance, this is repeated for the combined tensor *x*21. In both encoded tensors, the average over all non-padded sequence positions is computed for each embedding dimension, resulting in two vectors of length *d*_*att*_. These are combined into a single vector of length *d*_*att*_ by taking the position-wise maximum. This interaction feature vector is then turned into a prediction by a random feature expansion of a Gaussian process (RFEGP) layer, followed by a sigmoid function. All linear layers, including those in the encoders, are spectral-normalized. This, along with the RFEGP, serves to enhance the model’s uncertainty awareness while retaining accuracy. To test the influence of padding on the model performance, we implemented a version using unpadded per-token embeddings (“TUnA-unpadded”, for results, see Supplement). We also tested removing the spectral normalization and using a cross-attention mechanism in the encoders (“TUnA-no-spectral”, “TUnA-crossattention”). The family of TUnA adaptions is referred to as “TUnA-like”.

### 2.4 Distance map prediction

Further, we tested whether models with per-token embedding input can implicitly predict a contact map as a penultimate layer. This had first been suggested by Sledzieski et al. [2021] with their D-SCRIPT model. We performed a case study investigating the differences between real, experimentally determined distance maps and distance maps implicitly predicted by the models. We used distance maps instead of contact maps, as contact maps are binarized distance maps, indicating whether two positions are within a certain distance of each other [Badaczewska-Dawid et al., 2022]. To acquire the real distance maps, we searched the <u>Protein Data Bank (PDB)</u> [Berman et al., 2000] for protein complexes containing only the two proteins of the interactions of the gold-standard dataset, as other proteins, co-factors or ligands cannot be taken into consideration by the models. Complexes containing homomers were also ignored. We filtered out entries where structural information was available exclusively for small parts of the proteins, i.e., peptides stemming from the original protein. Still, the protein sequences from the PDB were mostly shorter than the sequences used as input for prediction. Additionally, many proteins had modifications at certain amino acids that were added in the experiment that generated the data. To account for both of these issues, local alignment was used to match the sequences from the predictions to those of the experimental data. We used only those predicted distance maps where the associated prediction was above the threshold of 0.9 to ensure the models’ confidence in the prediction. As a numerical value for similarity between experimentally determined and predicted distance maps, we computed the Pearson correlation of the two matrices.

## 3. Results

### 3.1. Hyperparameter optimization did not yield better parameter combinations than defaults

All models, apart from the Richoux-ESM-2 model, ran with 40 different hyperparameter configurations. Due to its considerably shorter runtime, approximately 250 different combinations were tested for the Richoux-ESM-2 model. The initial goal of the optimization was to identify the configuration that produces the best results. However, no parameter combination led to superior performances than the configurations using the original hyperparameter values (for the D-SCRIPT-ESM-2, Richoux-ESM-2, and TUnA) or the parameters that we chose originally (for 2d-Selfattention and 2d-Crossattention). Nonetheless, the optimization produced valuable insights into the impact of specific parameters on the performance, as well as into the robustness of the models themselves. For Richoux-ESM-2, the hyperparameters had minimal impact on the performance (Figure S7). For the models using an attention mechanism (2d-Selfattention, 2d-Crossattention, TUnA), the learning rate was always reported to be the most important parameter with high negative correlation (Figures S8-10). Finally, the 2d-Selfattention model favored max pooling, while the 2d-Crossattention model produced slightly better results with average pooling.

### 3.2. ESM-2 embeddings boost test accuracies of all models to around 0.65

For the performance evaluation of all models on the test set, we used the best-performing hyperparameter combinations (Table 1). The exact configurations can be found in the code of the associated GitHub repository. Of note, this resulted in all per-token models using the ESM-2 t33 embeddings and Richoux-ESM-2 using t36 embeddings. The RFC baseline with the best performance was RFC-40 on the t36 embeddings (Figure 2).

**Fig. 1:**
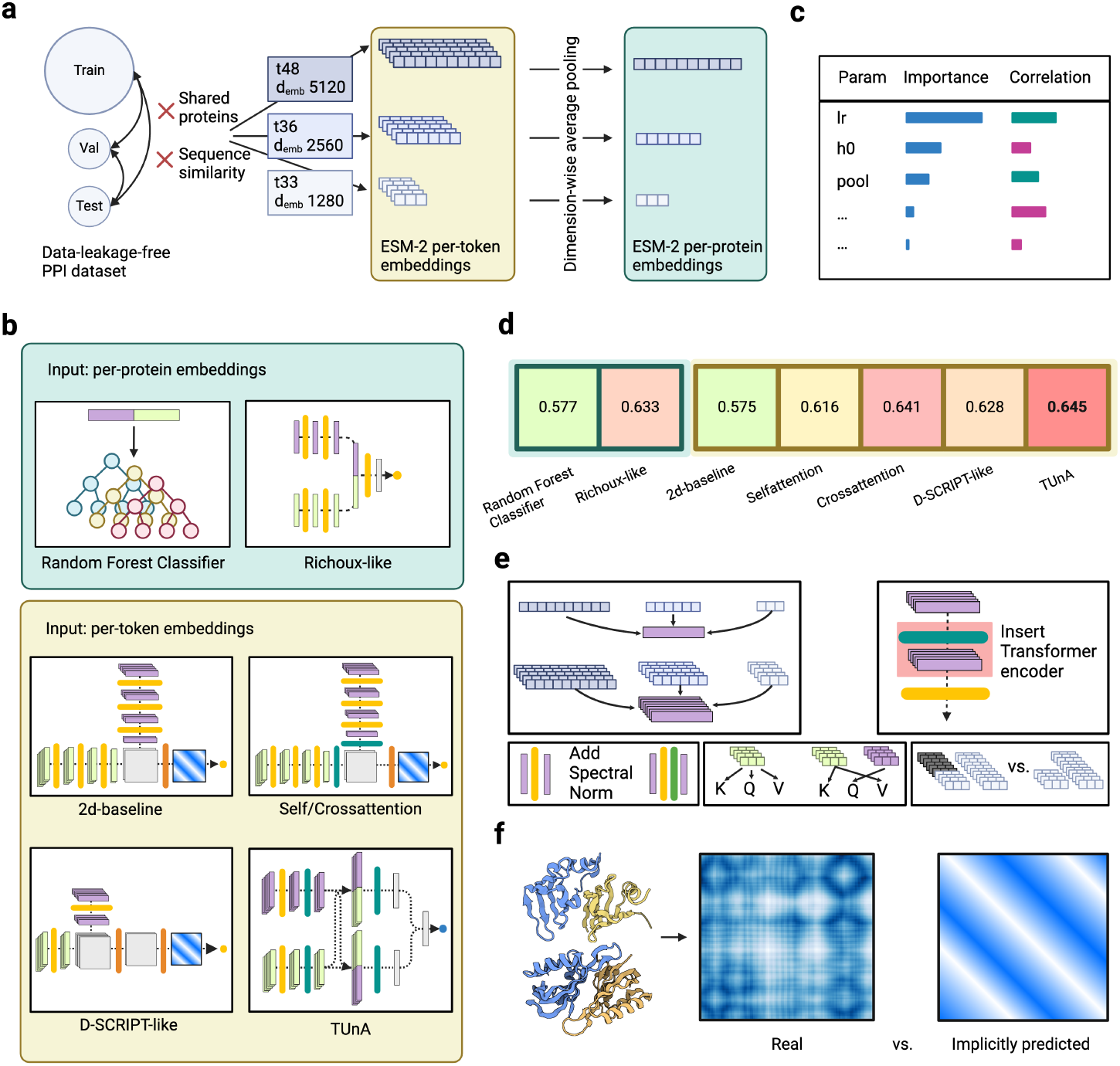
Overview of the analyses. **(a)** We computed ESM-2 embeddings of different sizes for the proteins of our data-leakage-free PPI dataset. The per-token embeddings have variable sizes depending on the protein length, while the per-protein embeddings have a fixed size by applying dimension-wise averaging. **(b)** We tested two models operating on the per-protein embeddings — a baseline random forest classifier and an adaptions the previously published Richoux model. Five models operated on the per-token embeddings: a 2d-baseline, the 2d-Selfattention and 2d-Crossattention models (which expanded the 2d-baseline through a Transformer encoder), and adaptations of the published models D-SCRIPT and TUnA. **(c)** Hyperparameter tuning gave us insight into the influence of each tunable parameter on the classification performance. **(d)** No model surpassed an accuracy of 0.65. The more advanced models had similar accuracies, leading us to believe that the information content of the ESM-2 embedding has more influence than the model architecture. Per-token models did not consistently outperform per-protein models. **(e)** We applied various modifications to test their influence: different embedding sizes, inserting a Transformer encoder into different positions, adding spectral normalization after the linear layers, self-vs. cross-attention, and removing the padding. **(f)** Finally, we compared the implicitly predicted distance maps of the 2d-baseline, 2d-Selfattention, 2d-Crossattention, and D-SCRIPT-ESM-2 to real distance maps computed from PDB structures. Created in https://BioRender.com

**Fig. 2:**
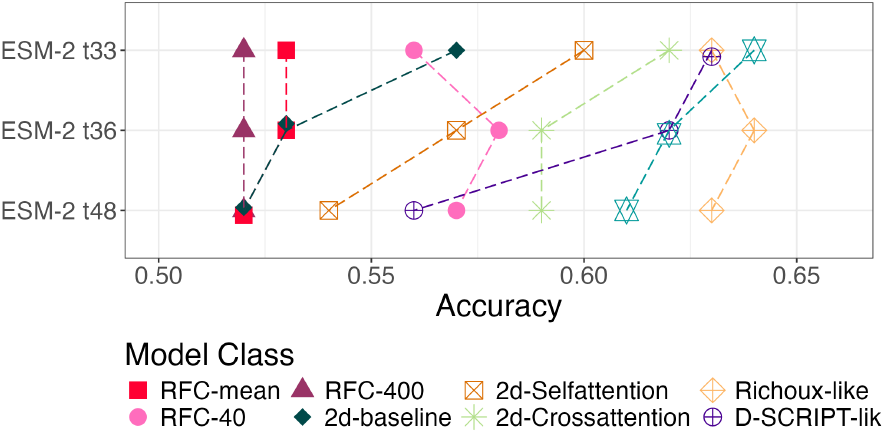
Validation performance with increasing ESM-2 embedding size. Most models perform best with the smaller t33 embedding.

TUnA achieves the highest accuracy, precision, and lowest BCE loss, closely followed by the 2d-Crossattention model. Notably, all models, except Richoux-ESM-2, exhibit much higher precision than recall, leading to reduced F1 scores. This imbalance is more pronounced in models with attention mechanisms, suggesting a conservative approach that minimizes false positives but sacrifices recall by missing true positives. In contrast, Richoux-ESM-2 achieves a better balance between precision and recall, improving its detection of true positives without significantly increasing false positives. None of the models deliver reliable PPI prediction, with accuracies not exceeding 0.65. All complex models fall within a narrow accuracy range of 0.61 to 0.65, regardless of their architectural differences. Using the ESM-2 embeddings, the adapted Richoux-ESM2 and D-SCRIPT-ESM2 now achieved accuracies of 0.633 and 0.628, demonstrating that the embeddings elevate models from near-random to near-state-of-the-art performance. Similarly, the RFC, previously achieving 0.53 accuracy, improves to 0.577, highlighting the embeddings’ information-rich content. This consistency suggests the performance is primarily driven by the ESM-2 embeddings rather than by the models themselves.

### 3.3. Per-token embeddings do not yield better results than averaged per-protein embeddings

An overview of the performance of all modifications can be found in Table S1. While the embeddings substantially enhance performance, the choice of embedding type (mean or per-token) has minimal impact. The baselines show virtually no difference. The more advanced models have no visible trend. While the best-performing models, TUnA and 2d-Crossattention, are per-token models, Richoux-ESM-2, which uses the mean embeddings, outperforms D-SCRIPT-ESM-2 and 2d-Selfattention. Further, the differences in performance are rather small overall. This finding is unexpected, as per-token embeddings should theoretically encode more critical interaction-specific information unavailable in mean embeddings. Regarding training, models with mean embeddings are significantly faster. Complex models employing per-token embeddings require approximately 2000 seconds per epoch, whereas the Richoux-ESM-2 model reduces this by a factor of 13 with comparable performance. RFC-40 achieves an additional order-of-magnitude speedup, though this excludes PCA runtime.

### 3.4. Models profit from smaller embeddings

Sledzieski et al. [2024] stated that using smaller input embeddings led to superior results in training their model. Although embedding size had little impact on Richoux models, others, such as TuNA, 2d-baseline as well as 2d-Self- and 2d-Crossattention showed increased performance on the t33 embeddings compared to the larger t36 and t48 embeddings (Figure 2). The similar performance of the different embedding sizes on the training set could indicate a relationship between larger input embeddings and the susceptibility to overfitting, possibly due to the increase in model parameters.

### 3.5. Only the 2d-baseline profits from attention

The 2d-Selfattention and 2d-Crossattention models differ from the 2d-baseline only by a transformer encoder layer, which is inserted between the embedding reduction and the combined representation of both sequences (*len*(*p*_1_) *× len*(*p*_2_) *×* 64). This modification resulted in notable performance improvements. 2d-Crossattention outperforms 2d-Selfattention, likely due to its ability to capture long-range protein interactions that complement the short-range relationships captured by convolution layers. Although the Transformer encoder nearly doubles runtime per epoch, the performance gains justify this trade-off.

To test the impact of the attention mechanism on published model architectures, we compared the regular D-SCRIPT-ESM-2 and Richoux-ESM-2 models to the versions utilizing an encoder (D-SCRIPT-ESM-2-encoder-post-reduction, Richoux-ESM-2-encoder-spectral). For Richoux-ESM-2-encoder-spectral, the attention mechanism resulted in a performance decrease (Figure 3). This decrease is likely due to an incompatibility of mean embeddings and the attention mechanism. In self-attention, the goal is to capture relationships in the input by modifying the embedding space of the related positions accordingly. By taking the average feature values of all sequence positions, the related positions enhanced by the self-attention will lose any meaning. Furthermore, the average feature values are altered by the attention mechanism, possibly making them partially unreadable or uninterpretable.

**Fig. 3:**
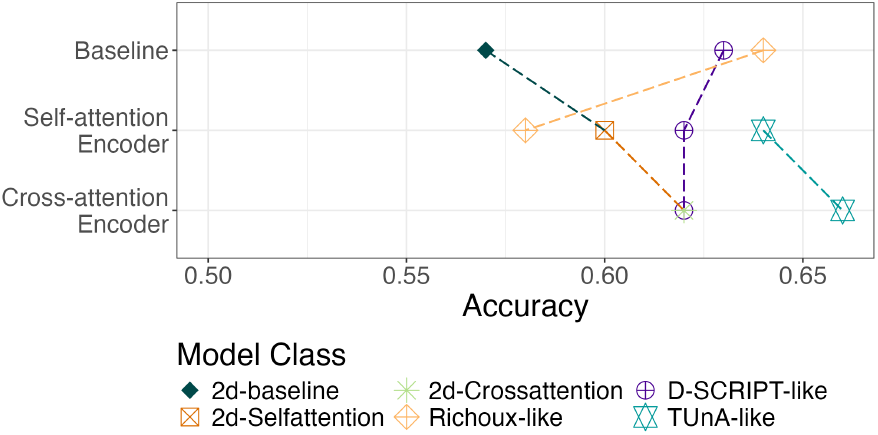
Validation performances before and after adding a self-/cross-attention encoder to the respective models.

For D-SCRIPT-ESM-2-encoder-post-reduction, the addition of the encoder had no notable influence on the model performance. This was surprising, given that the 2d-Selfattention and 2d-Crossattention models have a similar architecture as D-SCRIPT-ESM-2. One reason could be the performance barrier of 0.65 accuracy, which seems to be the information content of the ESM-2 embeddings. A model that can already harness the information contained in the embeddings and manages to approach the barrier by itself may not be able to benefit from the attention mechanism. A simple model that is not yet able to capture this information fully might be enhanced to do so through the attention mechanism.

As the 2d-Crossattention model outperforms the 2d-Selfattention model, we explored this aspect further by examining the influence of cross-attention versus self-attention on the D-SCRIPT-ESM-2-encoder-post-reduction and TUnA models (Figure 3). The D-SCRIPT model shows similar performance regardless of the type of attention mechanism while TUnA-crossattention is slightly better. The results of TUnA-crossattention also question the meaningfulness of the TUnA architecture, as the idea of having an intra-protein self-attention encoder that captures relationships inside each of the proteins and an inter-protein self-attention encoder that captures relationships between the two proteins is similar to adding cross-attention.

### 3.6. Attention-based models profit from spectral normalization

The TUnA publication argues that adding spectral normalization after linear layers increases model performance. Indeed, removing all spectral normalization layers in the TUnA model resulted in random predictions (Figure 4). Motivated by this behavior, we investigated the influence of adding spectral normalization to the other models. On Richoux-ESM-2, spectral normalization on the linear layers had no clear effect on the performance. On all attention-based models (2d-Selfattention, 2d-Crossattention, D-SCRIPT-ESM-2-encoder-post-reduction, and Richoux-ESM-2-encoder), the absence of spectral normalization also resulted in random predictions (Figure 4).

**Fig. 4:**
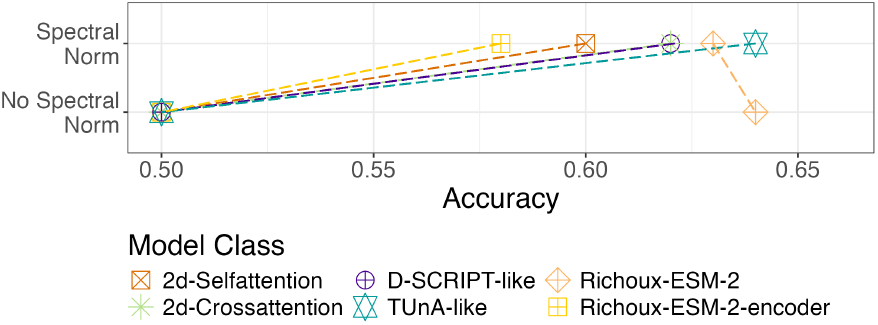
Validation performance before and after removing spectral normalization after the linear layers for the models including an encoder and Richoux-ESM-2.

While investigating possible causes of this, we discovered that models using any non-spectral normalized attention mechanisms stagnated during training. Irrespective of the input, the model went back and forth between either predicting all interactions as positive, with values slightly above 0.5, or negative, with values slightly below 0.5. We can only speculate about the reasons for this behavior. It could be that without spectral normalization, the gradients in the attention mechanism exponentially grow or decay, leading to nonsensical output. Interestingly, the TUnA-no-spectral (Figure S11) even learns the data regularly for the first few epochs before also dropping to random performance. Accordingly, all comparisons of models using attention mechanisms were done using spectral normalization.

### 3.7. Attention should be applied after reducing embedding size

We explored the influence of reducing the embedding size via linear layers prior to the attention mechanism or encoder. In all tested models, reducing embedding size leads to superior performance on the validation dataset (Figure 5). D-SCRIPT-ESM-2-encoder-pre-reduction had near-random performance scores. We hypothesize that not reducing the embedding dimension may have a similar effect as using a larger embedding, which, as stated before, may be the cause for increased overfitting, explaining the performance of the 2d-Selfattention model. The diminished performance of both 2d-Crossattention-encoder-pre-reduction and D-SCRIPT-ESM-2-encoder-pre-reduction on the training and validation sets may also hint at the negative impact of using many input parameters for the attention mechanism, as it could potentially obscure critical positions among the noise.

**Fig. 5:**
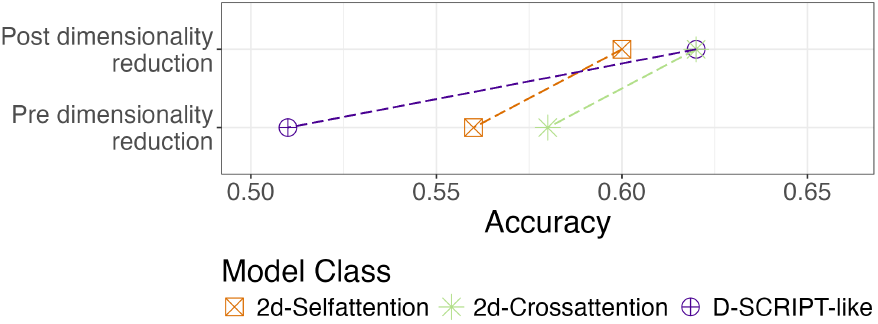
Validation performance for inserting the encoder before or after dimensionality reduction via linear layers.

### 3.8. Distance maps cannot be predicted implicitly

Of the tested models, D-SCRIPT-ESM-2, the 2d-baseline, 2d-Selfattention, and 2d-Crossattention internally form a matrix of size *len*(*p*_1_) *× len*(*p*_2_). As D-SCRIPT [Sledzieski et al., 2021] treats this as implicit contact map prediction, we investigated the similarities of this matrix to the real, experimentally determined contact maps. For this case study, we used all protein pairs of the dataset, regardless of the split they appeared in, as the models cannot overfit the distance maps during training because they received no structural information. Matching the confident predictions (*>* 0.9) with the PDB entries resulted in a total of 84 interactions across all models. Of those, 54 were filtered out due to them containing other cofactors, ligands, or homomers, and another 19 were removed because the protein sequences in the experimental data were too short. This left 3 interactions for D-SCRIPT-ESM-2, 1 for 2d-Crossattention, 7 for 2d-Selfattention, and none for the 2d-baseline (Table S2).

In the predicted distance maps from D-SCRIPT-ESM-2, certain structures are formed, which is especially visible in Figure 6. In Figure S12, some vertical lines from the real distance map can be found in the prediction, indicating that the model identified some of the important positions of the Q8NAV1 protein. In the 1B34 complex (Figure S13), there are several clusters of high values in the bottom left and clusters of low values in the top right corner. While these clusters do not align with any clear features of the real distance map, they do exhibit a slight negative correlation.

**Fig. 6:**
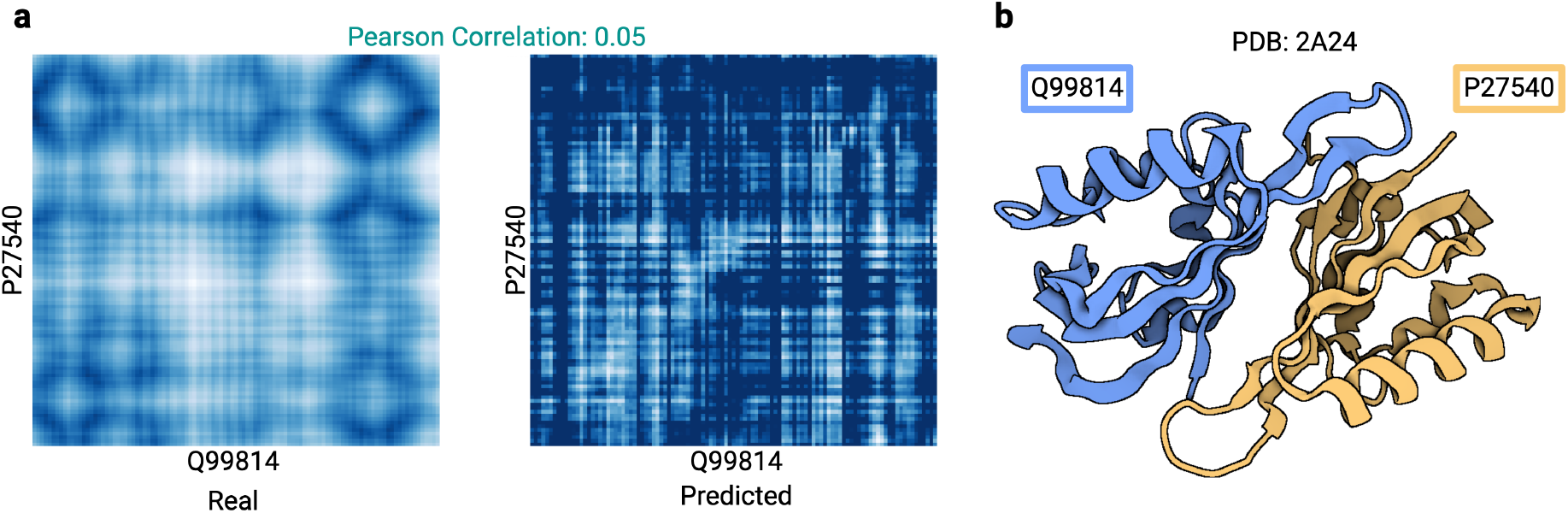
**(a)** Comparison of the real and predicted distance map of the D-SCRIPT-ESM-2 model for the 2A24 complex. White indicates low values (i.e., contact in the distance map), dark blue high values. **(b)** PDB complex of 2A24, created in https://BioRender.com

Like D-SCRIPT-ESM-2, the 2d-Selfattention model also confidently predicted the interaction of complex 1B34 (Figure S14). In contrast to D-SCRIPT-ESM-2, no clear, distinct clusters or features are formed. Rather, some horizontal lines can be identified in both distance maps, hinting at the successful identification of some important positions of P63214. While both models achieve the same, although inverse, correlation, the 2d-Selfattention prediction resembles a seemingly structureless grid rather than an actual distance map. The same grid can be observed in the predictions for 2V8S for both 2d-Self- and 2d-Crossattention. Interestingly, 2d-Selfattention more clearly detects the vertical lines (Figure S14), unlike the 2d-Crossattention model, which more distinctly recognizes the horizontal features (Figure S16). Similar results can be observed for the remaining maps predicted by 2d-Selfattention (Figures S17–S21).

Ultimately, the observed low correlations and the notable disparity between the real and predicted maps show that no model is capable of indirectly predicting distance maps or even detecting more complex structural features. This was to be expected, considering that the models receive only sequential data and that the task of predicting 3D structure from sequence was only partially solved by AlphaFold [Jumper et al., 2021], which is considerably more complex. However, the models were able to identify some singular, important positions for the interaction.

## 4. Discussion

In this study, we comprehensively investigated the efficacy in PPI prediction of various deep learning models, including Richoux-ESM-2, D-SCRIPT-like, and TUnA-like models, as well as 2d-Selfattention, and 2d-Crossattention models. Hyperparameter optimization across 40 configurations had little impact, and all models struggled to surpass an accuracy of 0.65, implying that the reported performance might be dictated more by the embeddings than the architecture of the individual models. This is further supported by the literature, where similar results are reported. In terms of speed, models utilizing averaged per-protein embeddings proved faster than their per-token counterparts, while producing similar results. We also tested several modifications of the models. For the relatively simple 2d-Crossattention and 2d-Selfattention models, adding a Transformer encoder with an attention mechanism led to a notable increase in performance compared to the 2d-baseline. In contrast, performances did not improve for more complex models, which reached test accuracies close to 0.65 already without Transformer encoders. This further supports our hypothesis that this value constitutes a barrier that is unlikely to be surpassed by purely sequence-based models.

In comparing real distance maps with those predicted by the models, we tested the hypothesis of implicitly predicting contact maps as suggested in the D-SCRIPT paper [Sledzieski et al., 2021]. While the D-SCRIPT-ESM-2 model produced some recognizable structures in these maps, they showed little to no relation to the experimentally determined distance maps. Similar results were observed for the 2d-Selfattention and 2d-Crossattention models. These findings highlight the broader challenge of accurately predicting structural data in pairwise PPIs, given the limited availability of such data.

Despite these highly relevant findings, our study is subject to at least two limitations. The first limitation was computational power. As mentioned in the Methods section, many compromises were made due to computational limitations, such as restricting the input in embedding size and sequence length, as well as an inexhaustive optimization of model hyperparameters. Secondly, the availability of PPI data for training, validation, and testing also poses an issue. While our gold-standard dataset is an improvement compared to previous PPI datasets, many top-performing ML models, such as AlphaFold [Jumper et al., 2021], are trained on substantially larger datasets.

When screening PDB for data on protein complexes for our comparison between experimentally determined and implicitly predicted distance maps, we identified only very few suitable complexes that contain only two proteins. The PDB reports 2836 pairwise protein interactions without any cofactors, ligands or other proteins, while it reports 46 849 entries with at least one additional subunit. Furthermore, some interactions described as involving only two proteins may actually consist of homomers. This points to a general limitation of existing sequence-based PPI prediction models: While existing models are designed to predict PPIs consisting of only two proteins, most complexes formed in permanent interactions are comprised of many more proteins, other cofactors or ligands, which can not be captured in pairwise interactions but may be essential for the interaction.

Another key challenge lies in the choice of protein embeddings, as both mean and per-token approaches present significant limitations for sequence-based PPI prediction. Mean embeddings lack positional information, making them unsuitable for capturing structural details such as interacting amino acids, especially when all positions are weighted equally during the average pooling. In contrast, per-token embeddings retain positional data but introduce variability in sequence length, requiring computationally expensive padding or alternative handling. Padding, while standard, can inflate computational costs and lead to uneven training due to sequence length distributions. Moreover, models trained on padded sequences may fail to process proteins longer than the training limit. Unpadded embeddings avoid these issues but restrict sequence length reductions to basic operations, limiting flexibility. Per-token embeddings may perform well for position-specific tasks (e.g., predicting the effects of single nucleotide polymorphisms), but their application to PPI is constrained by the need for experimental identification of contact positions in unseen proteins.

Given these limitations, we suggest the following directions for future work: First, enhancing the embeddings with non-sequential data might boost the performance. Some approaches integrating natural language embeddings of protein functionality descriptions have already been proposed. Ko et al. [2024b] even reach an accuracy of 0.68 on our dataset by fusing the embeddings of several of those approaches. Further, incorporating structural information could address the limitations of sequence-based PPI prediction, although the current scarcity of suitable data may not allow for this yet. Another option is to predict protein structure as an intermediate step, leveraging tools like AlphaFold, which provide high-quality structural predictions. These predictions could enable the creation of larger artificial datasets, blending structural insights with sequence-based methods. Predicting contact regions or specific interaction sites could further refine inputs, reducing noise and enhancing the utility of per-token embeddings.

## 5. Code availability

All code for models and execution of the models is available at https://github.com/daisybio/PPI_prediction_study. Python version 3.8.18 and PyTorch (Paszke et al. [2019]) version 2.1.1 were used for this study. The environment containing the versions of all other packages used can be found on the GitHub repository.

## 7. Author contributions statement

J.B. and M.L. designed and conceived this study and supervised the work. T.R. implemented the test protocol, carried out the analyses, and drafted the manuscript. A.H. and D.B.B. gave critical feedback and made suggestions for improvements. All authors contributed to and approved the final manuscript.

## 8. Acknowledgments

T.R., A.H., and D.B.B. were supported by the German Federal Ministry of Education and Research (BMBF) within the framework of the CompLS funding concept [031L0309A (NetMap)]. M.L. and D.B.B. were supported by the Klaus Tschira Stiftung [00.003.2024]. J.B. and M.L. were supported by the German Federal Ministry of Education and Research (BMBF) within the framework of the CompLS funding concept [031L0305A (DROP2AI)]. Funded by the Deutsche Forschungsgemeinschaft (DFG, German Research Foundation) [422216132].

