## Supplemental Figures and Tables for "Deep learning models for unbiased sequence-based PPI prediction plateau at an accuracy of 0.65"

Supplementary information

### 1 Additional results: padding is unnecessary for TUnA

For the TUnA model, we also examined the impact of removing the padding from the input embeddings on the performance. As the TUnA model never uses any modules with trainable parameters to reduce the sequence dimension as well as mask padded positions in the attention mechanisms, we expected there to be no difference between using unpadded and padded inputs. This suspicion was confirmed by the test (Figure S11).

### 2 Supplementary figures and tables

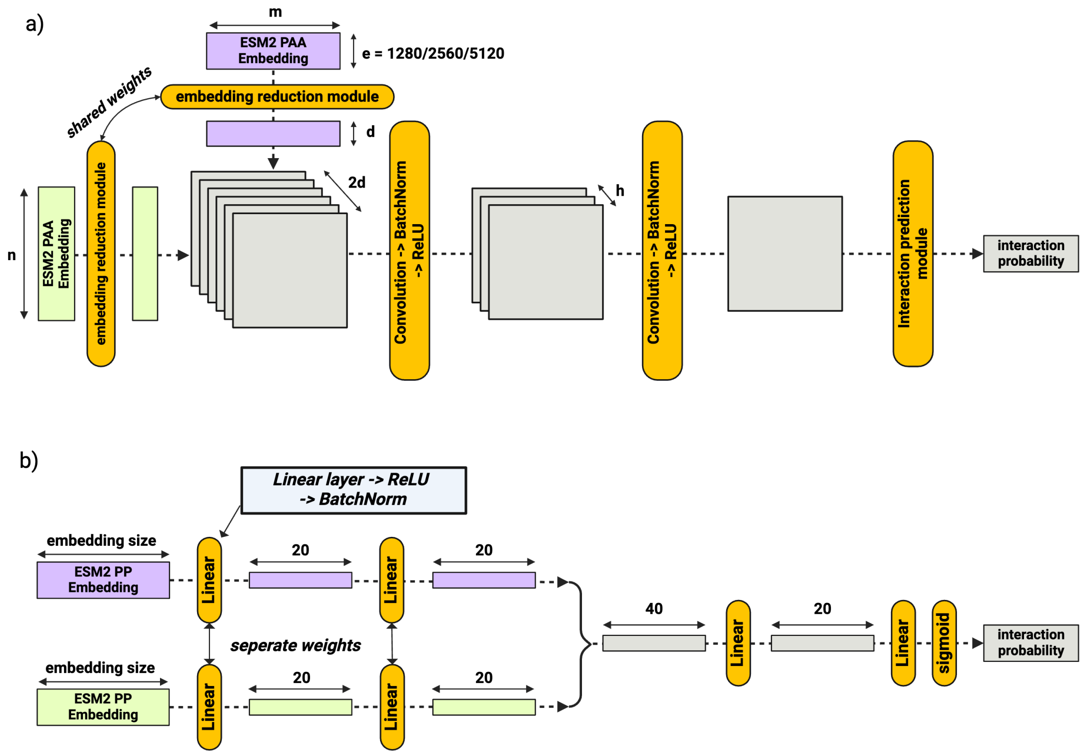


Figure S1: Visual representation of used Models. Rectangles represent the data, dimensions shown by arrows. Curved rectangles represent the modules used to change the data; a) D-SCRIPT-like; b) Richoux-like. Created in https://BioRender.com

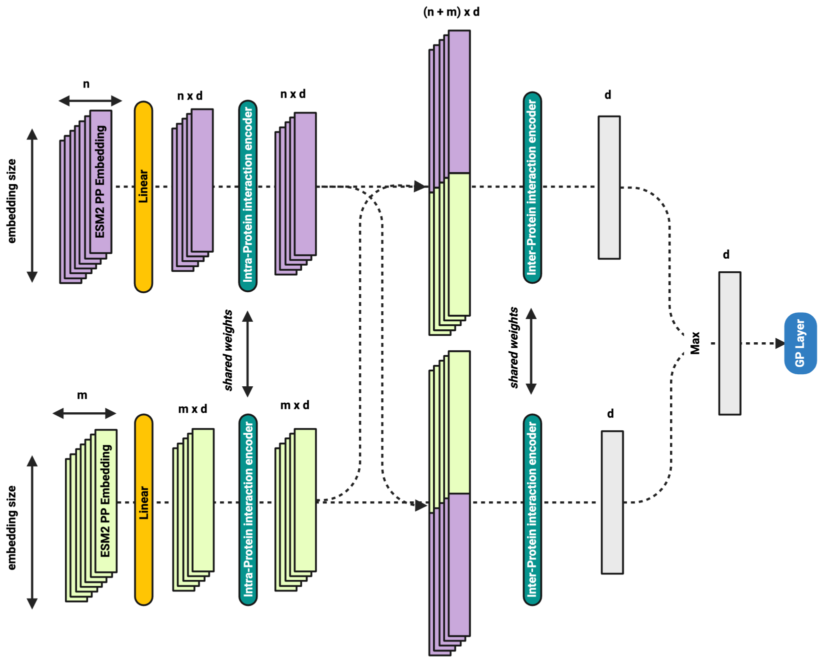


Figure S2: Visual representation of TUnA model architecture. Created in https://BioRender.com

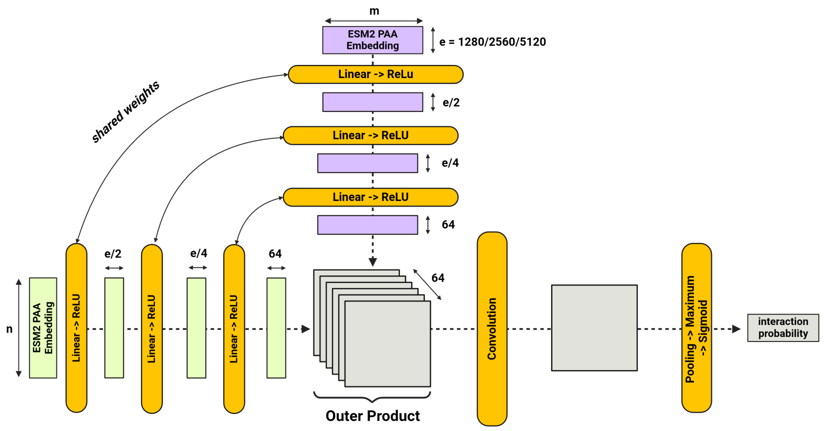


Figure S3: Visual representation of 2d-baseline. Rectangles represent the data, dimensions shown by arrows. Curved rectangles represent the modules used to change the data. Created in https://BioRender.com

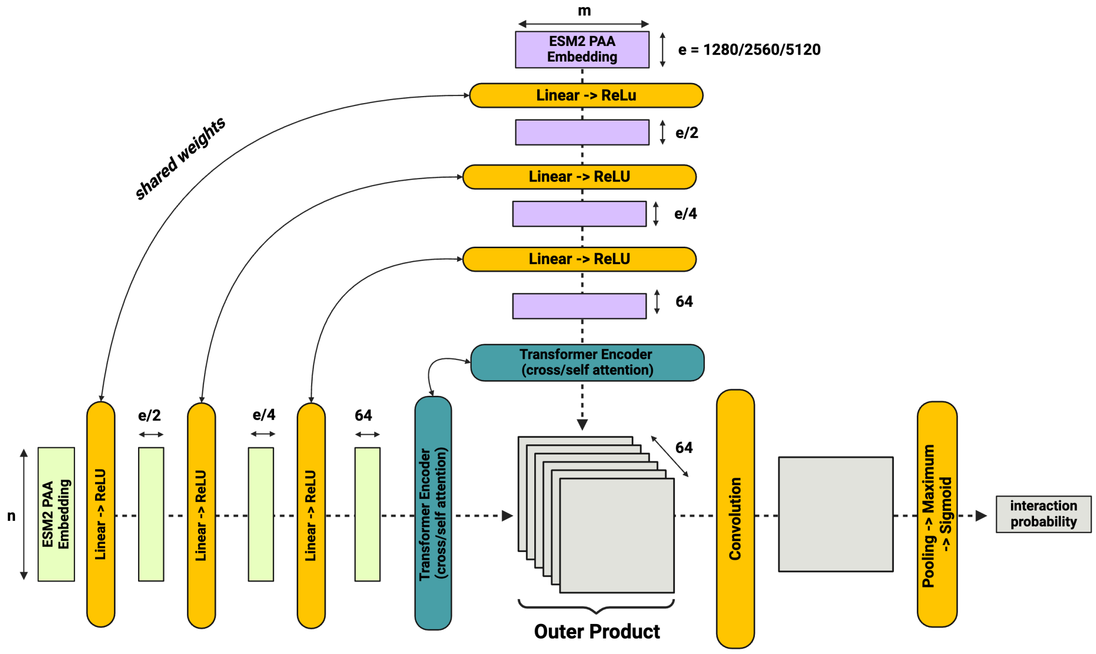


Figure S4: Visual representation of the attention models. Rectangles represent the data, dimensions shown by arrows. Curved rectangles represent the modules used to change the data. Created in https://BioRender.com

Table S1: Overview of all tested configurations.

| Model | Accuracy | Model Class | Modification |
|---|---|---|---|
| RFC-40 t33 | 0.56 | RFC | Base |
| RFC-40 t36 | 0.58 | RFC | Larger Embedding |
| RFC-40 t48 | 0.57 | RFC | Larger Embedding |
| RFC-400 t33 | 0.52 | RFC | Base |
| RFC-400 t36 | 0.52 | RFC | Larger Embedding |
| RFC-400 t48 | 0.52 | RFC | Larger Embedding |
| RFC-mean t33 | 0.53 | RFC | Base |
| RFC-mean t36 | 0.53 | RFC | Larger Embedding |
| RFC-mean t48 | 0.52 | RFC | Larger Embedding |
| 2d-baseline t33 | 0.57 | 2d-baseline | Base |
| 2d-baseline t36 | 0.53 | 2d-baseline | Larger Embedding |
| 2d-baseline t48 | 0.52 | 2d-baseline | Larger Embedding |
| 2d-Selfattention t33 | 0.60 | 2d-Selfattention | Base |
| 2d-Selfattention t36 | 0.57 | 2d-Selfattention | Larger Embedding |
| 2d-Selfattention t48 | 0.54 | 2d-Selfattention | Larger Embedding |
| 2d-Selfattention-no-spectral | 0.50 | 2d-Selfattention | No Spec. Norm |
| 2d-Selfattention-encoder-pre-reduction | 0.56 | 2d-Selfattention | Encoder pre reduction |
| 2d-Crossattention t33 | 0.62 | 2d-Crossattention | Base |
| 2d-Crossattention t36 | 0.59 | 2d-Crossattention | Larger Embedding |
| 2d-Crossattention t48 | 0.59 | 2d-Crossattention | Larger Embedding |
| 2d-Crossattention-encoder-pre-reduction | 0.58 | 2d-Crossattention | Encoder pre reduction |
| 2d-Crossattention-no-spectral | 0.50 | 2d-Crossattention | No Spec. Norm |
| Richoux-ESM-2 t33 | 0.63 | Richoux-like | Base |
| Richoux-ESM-2 t36 | 0.64 | Richoux-like | Larger Embedding |
| Richoux-ESM-2 t48 | 0.63 | Richoux-like | Larger Embedding |
| Richoux-ESM-2-spectral | 0.63 | Richoux-like | Spec. Norm |
| Richoux-ESM-2-encoder-no-spectral | 0.50 | Richoux-like | No Spec. Norm |
| Richoux-ESM-2-encoder-spectral | 0.58 | Richoux-like | Encoder pre reduction |
| D-SCRIPT-ESM-2 t33 | 0.63 | D-SCRIPT-like | Base |
| D-SCRIPT-ESM-2 t36 | 0.62 | D-SCRIPT-like | Larger Embedding |
| D-SCRIPT-ESM-2 t48 | 0.56 | D-SCRIPT-like | Larger Embedding |
| D-SCRIPT-ESM-2-encoder-pre-reduction | 0.51 | D-SCRIPT-like | Encoder pre reduction |
| D-SCRIPT-ESM-2-encoder-pre-reduction-no-spectral | 0.50 | D-SCRIPT-like | No Spec. Norm |
| D-SCRIPT-ESM-2-encoder-post-reduction | 0.62 | D-SCRIPT-like | Encoder post reduction |
| D-SCRIPT-ESM-2-encoder-crossattention | 0.62 | D-SCRIPT-like | Cross-attention |
| TUnA t33 | 0.64 | TUnA-like | Base |
| TUnA t36 | 0.62 | TUnA-like | Larger Embedding |
| TUnA t48 | 0.61 | TUnA-like | Larger Embedding |
| TUnA-crossattention | 0.66 | TUnA-like | Cross-attention |
| TUnA-unpadded | 0.65 | TUnA-like | No padding |
| TUnA-no-spectral | 0.50 | TUnA-like | No Spec. Norm |

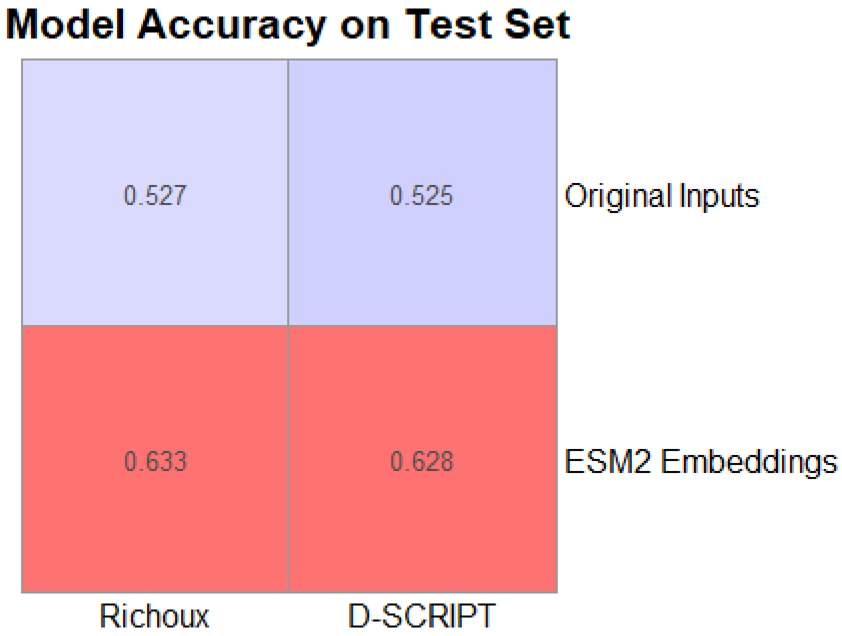


Figure S5: Comparison of using the original inputs versus ESM2 embeddings for the Richoux and D-SCRIPT models

Table S2: Amount of filtered PPIs due to additional cofactors or ligands, homomers or short length; Interactions is the total number of entries in the PDB with exactly two distinct proteins that the models predicted confidently (> 0.9); Chains is the number of entries filtered out due to additional chains, i.e., cofactors, ligands or homomers; Length describes the number of entries filtered out due to at least one protein sequence being too short; Total Skipped is the total amount of skipped entries per model (does not account for duplicate entries); Percentage shows the percentage of filtered entries.

| Model | Interactions | Chains | Length | Total Skipped | Percentage |
|---|---|---|---|---|---|
| DSCRIPT-like | 21 | 13 | 5 | 18 | 86% |
| Crossattention | 12 | 8 | 3 | 11 | 92% |
| Selfattention | 32 | 18 | 7 | 25 | 78% |
| 2d-baseline | 19 | 15 | 4 | 19 | 100% |
| Total | 84 | 54 | 19 | 73 | 87% |

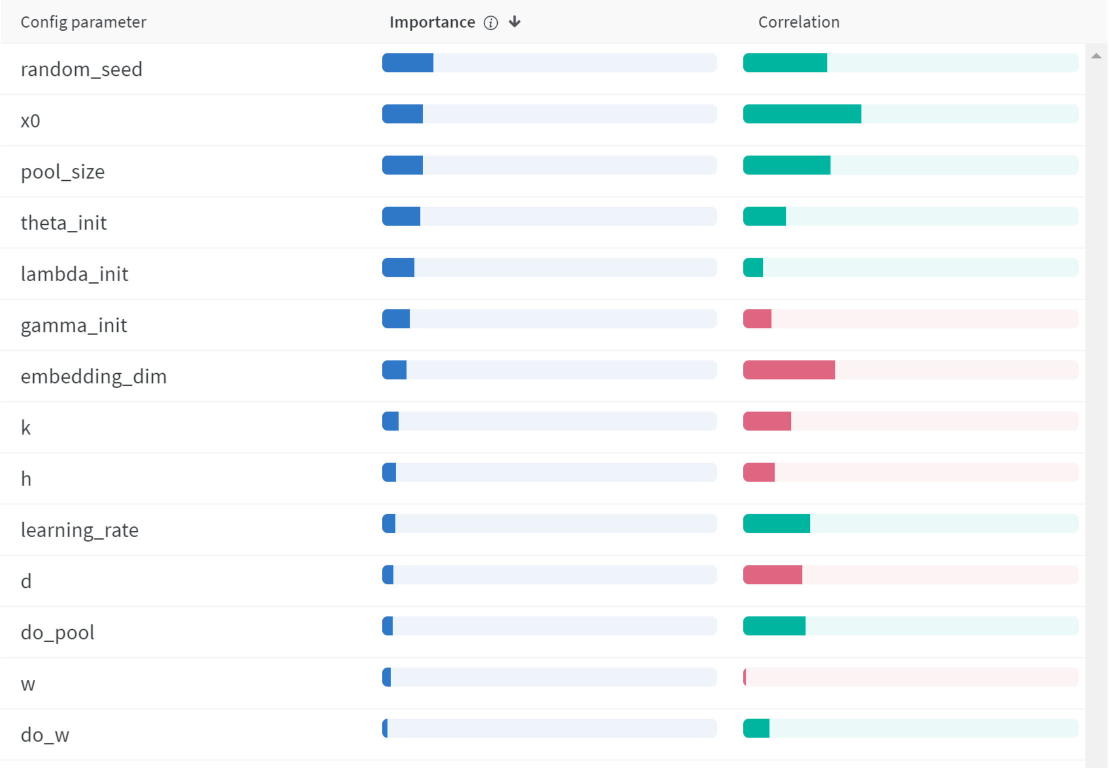


Figure S6: Importance and Correlation of hyperparameters of the DSCRIPT-like model. “d” and “h” are used to reduce embedding size in the linear layer and convolution; “pool_size” and “w” control the kernel size of pooling and convolution, respectively; “theta_init” and “lambda_init” are used in the weighing of the contact map; “k”, “gamma_init” and “x0” are parameters for the custom activation function; “do_pool” and “do_w” control whether the pooling and weighing will be performed. Created with wandb.

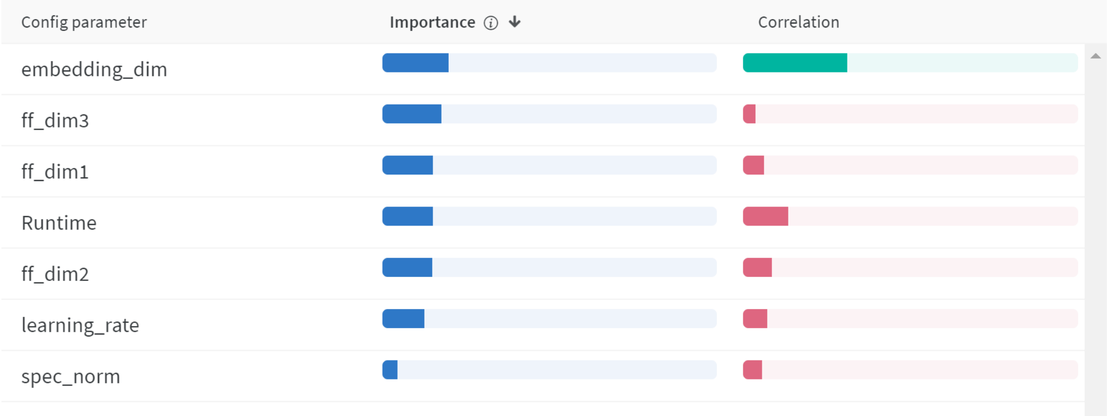


Figure S7: Importance and Correlation of hyperparameters of the Richoux-like model, Created with wandb.

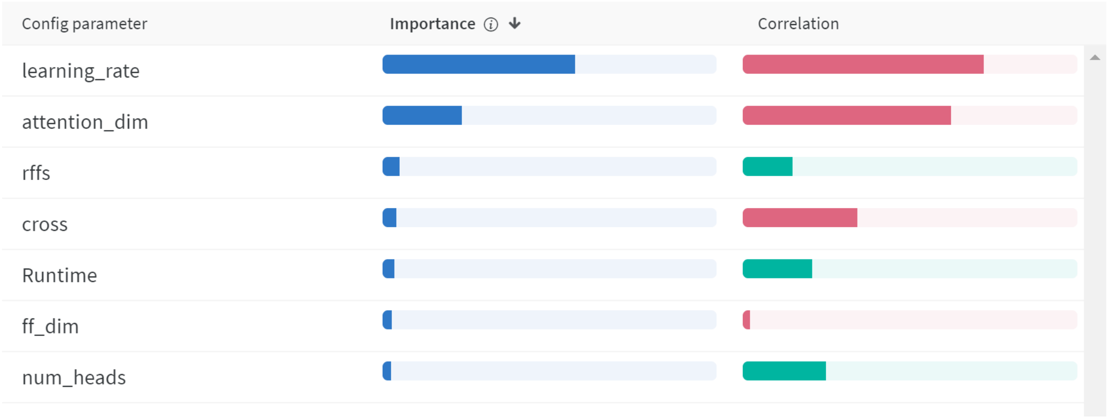


Figure S8: Importance and Correlation of hyperparameters of the TUnA model, Created with wandb.

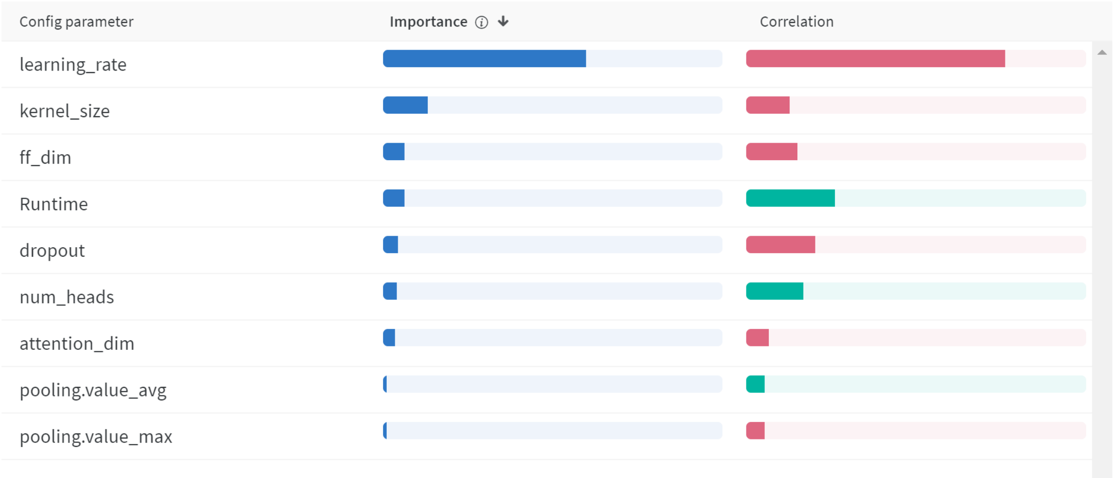


Figure S9: Importance and Correlation of hyperparameters of the Crossattention model, Created with wandb.

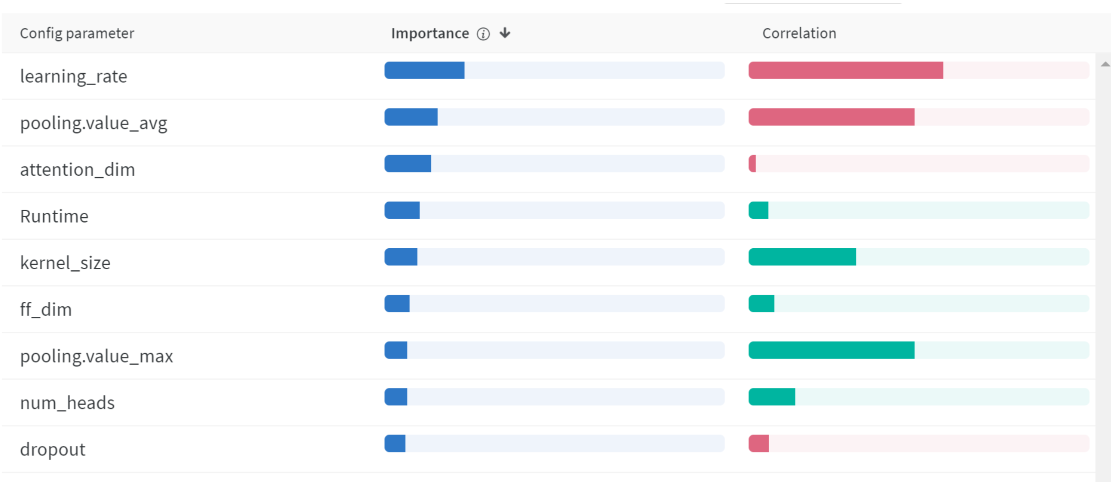


Figure S10: Importance and Correlation of hyperparameters of the Selfattention model, Created with wandb.

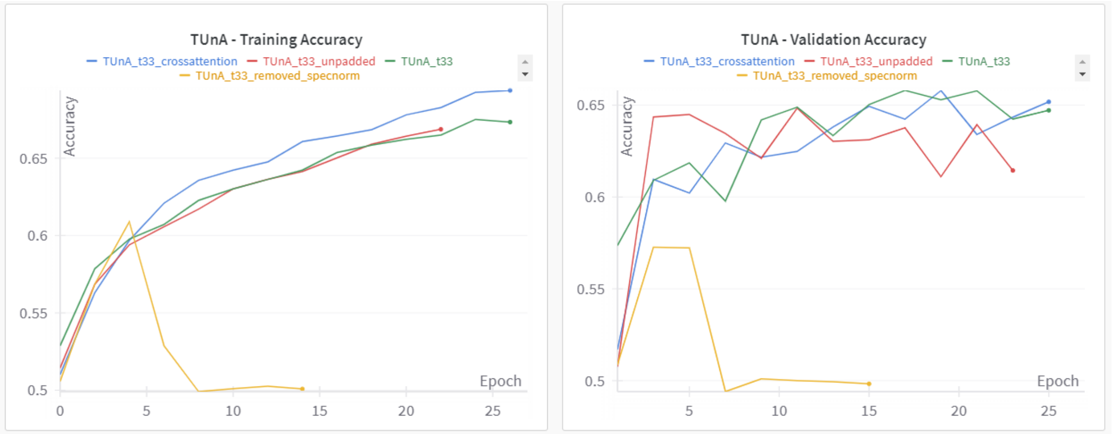


Figure S11: Comparison of TUnA models. Unmodified TUnA (green), TUnA with unpadded input (red), TUnA with cross attention in the encoders (blue), TUnA without spectral normalization (yellow). Created with wandb.

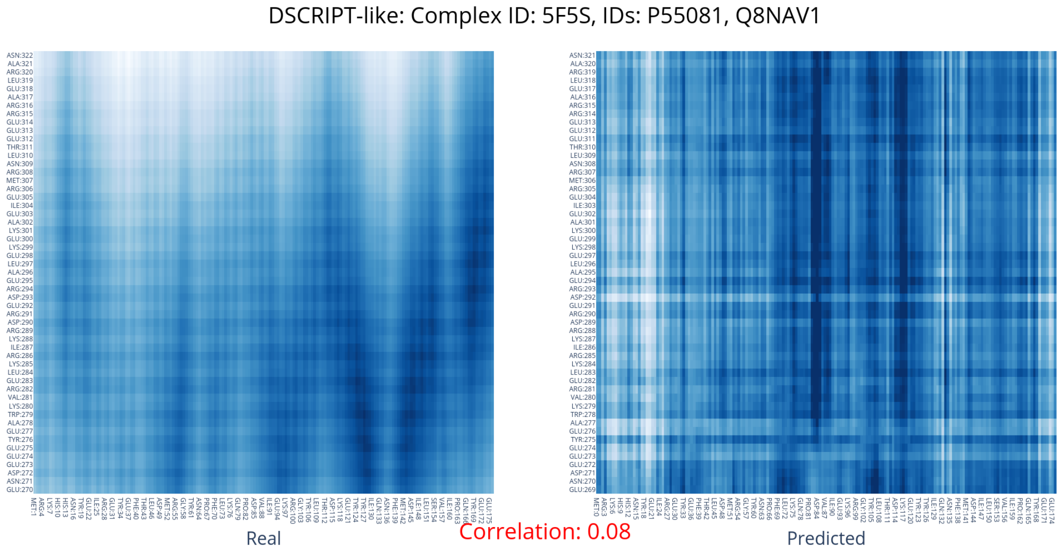


Figure S12: Comparison of the real and predicted distance map of the DSCRIPT-like model for the 5F5S complex. White indicates low values (i.e., contact in the distance map), dark blue high values.

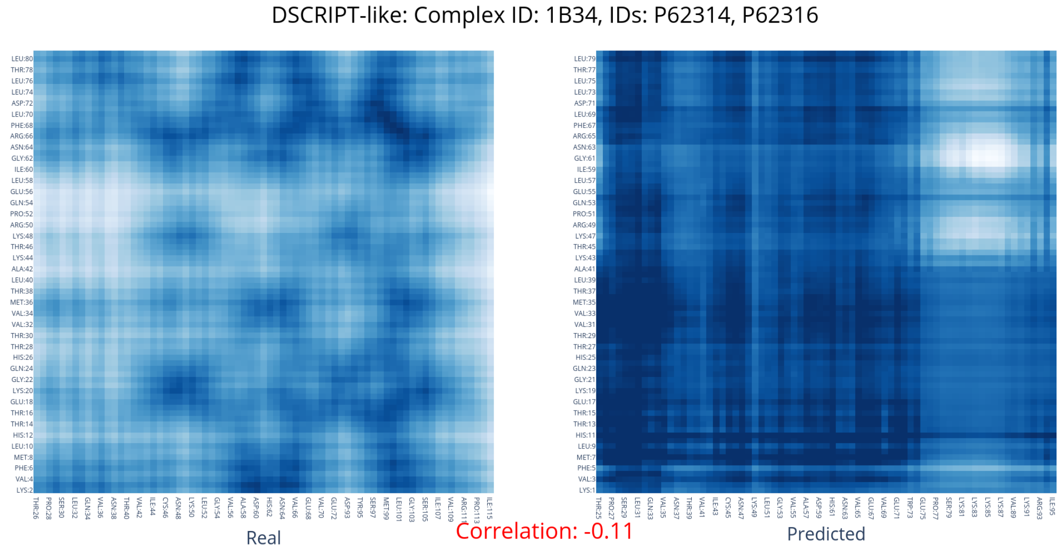


Figure S13: Comparison of the real and predicted distance map of the DSCRIPT-like model for the 1B34 complex. White indicates low values (i.e., contact in the distance map), dark blue high values.

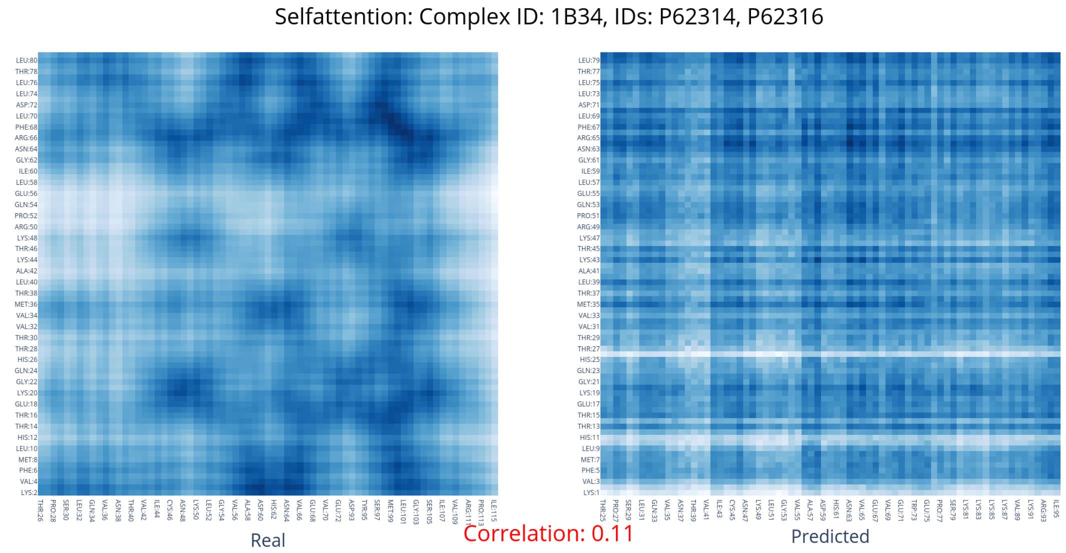


Figure S14: Comparison of the real and predicted distance map of the Selfattention model for the 1B34 complex. White indicates low values (i.e., contact in the distance map), dark blue high values.

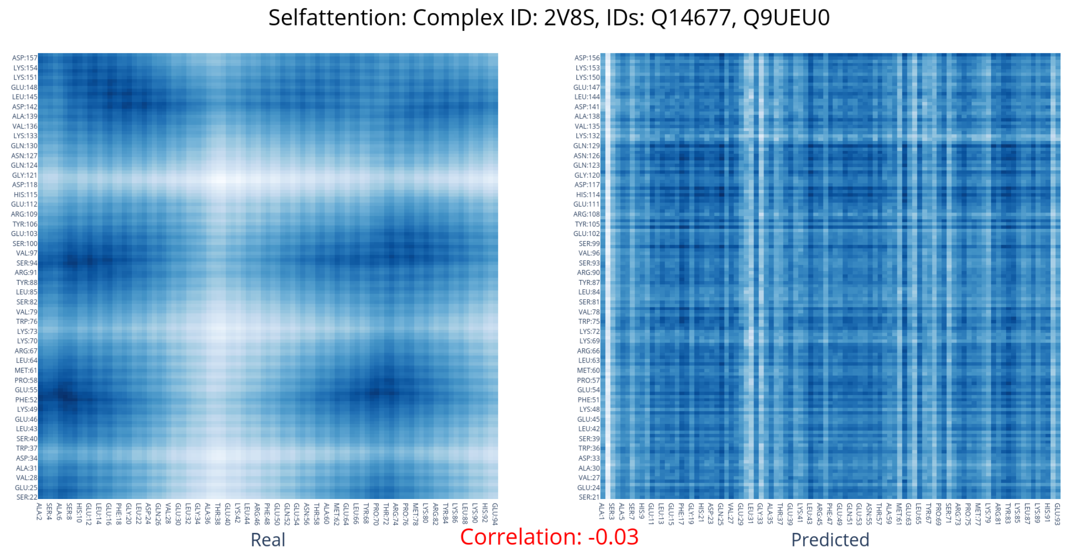


Figure S15: Comparison of the real and predicted distance map for the 2V8S: Selfattention model. White indicates low values (i.e., contact in the distance map), dark blue high values.

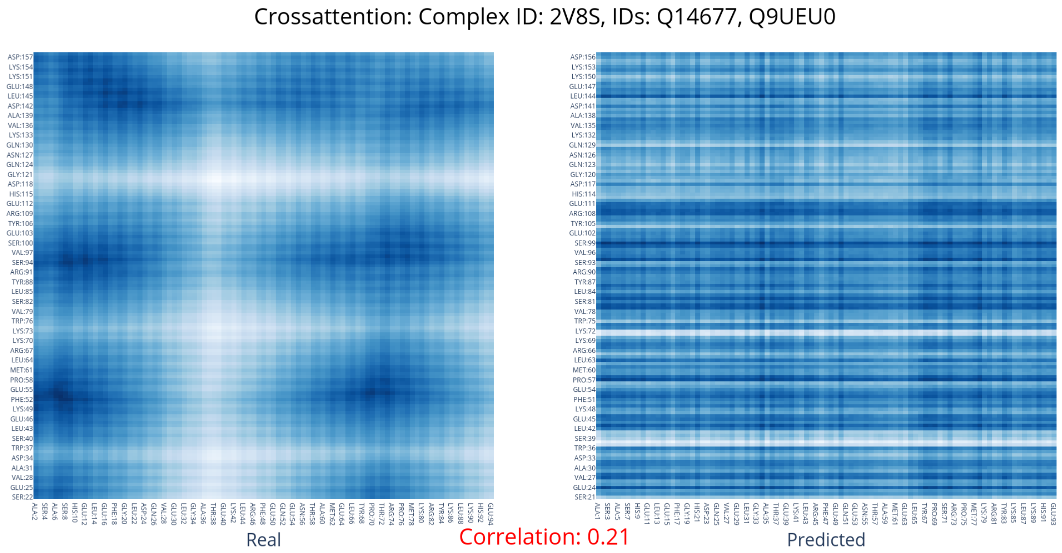


Figure S16: Comparison of the real and predicted distance map for the 2V8S: Crossattention model. White indicates low values (i.e., contact in the distance map), dark blue high values.

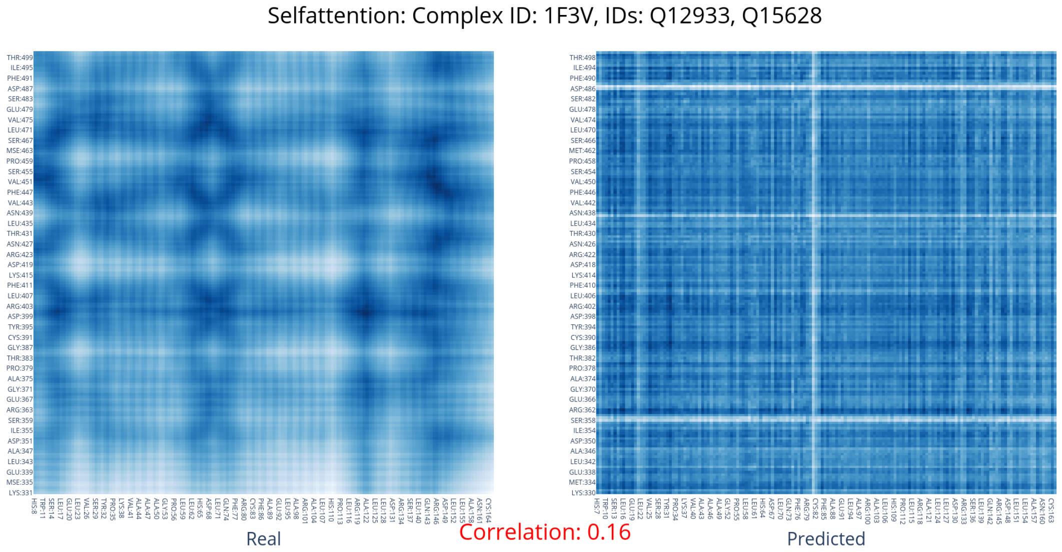


Figure S17: Comparison of the real and predicted distance map of the Selfattention model for the 1F3V complex. White indicates low values (i.e., contact in the distance map), dark blue high values.

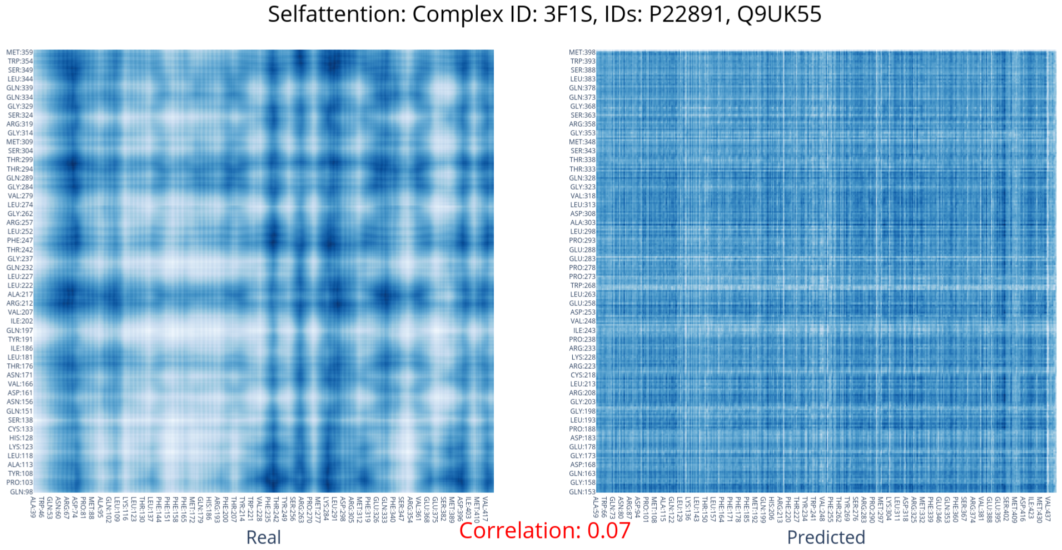


Figure S18: Comparison of the real and predicted distance map of the Selfattention model for the 3F1S complex. White indicates low values (i.e., contact in the distance map), dark blue high values.

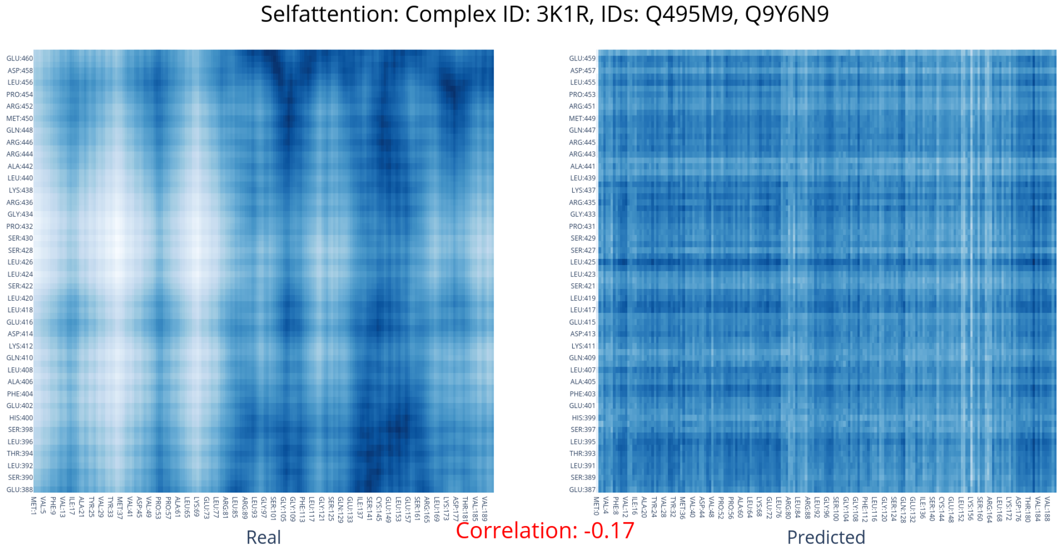


Figure S19: Comparison of the real and predicted distance map of the Selfattention model for the 3K1R complex. White indicates low values (i.e., contact in the distance map), dark blue high values.

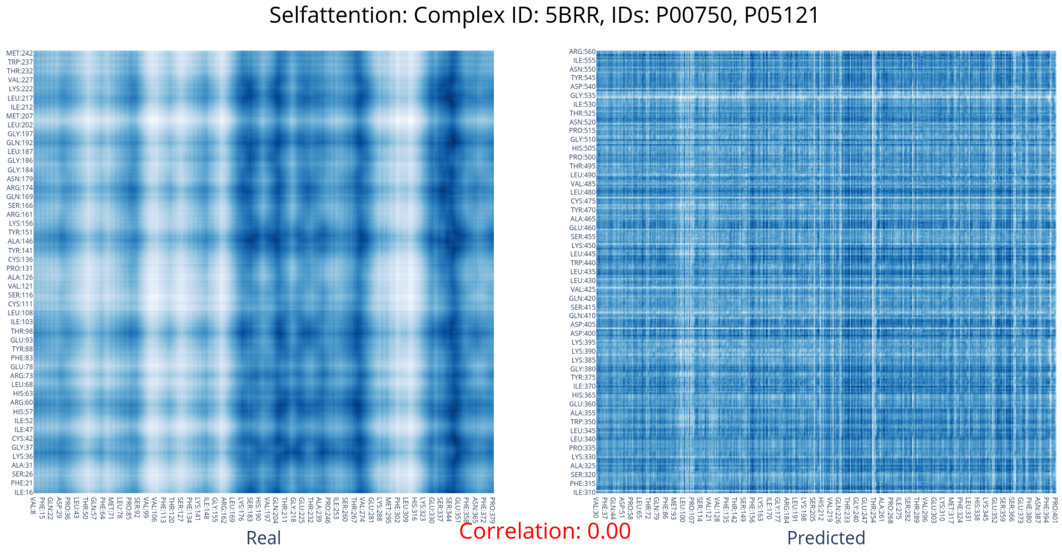


Figure S20: Comparison of the real and predicted distance map of the Selfattention model for the 5BRR complex. White indicates low values (i.e., contact in the distance map), dark blue high values.

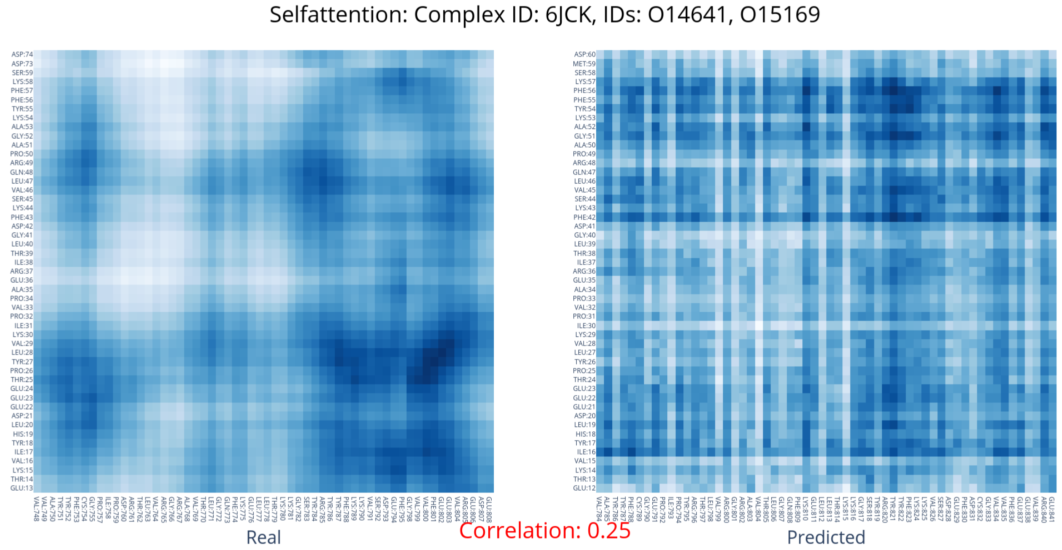


Figure S21: Comparison of the real and predicted distance map of the Selfattention model for the 6JCK complex. White indicates low values (i.e., contact in the distance map), dark blue high values.
